# Foodscapes for Salmon and Other Mobile Consumers in River Networks

**DOI:** 10.1101/2023.08.30.555604

**Authors:** Gabriel J. Rossi, J. Ryan Bellmore, Jonathan B. Armstrong, Carson Jeffres, Sean M. Naman, Stephanie M. Carlson, Theodore E. Grantham, Matthew J. Kaylor, Seth White, Jacob Katz, Mary E. Power

## Abstract

Mobile consumers track fluctuating resources across heterogeneous landscapes to grow and survive. In river networks, the abundance and accessibility of food and the costs of foraging vary between habitats and through time, providing a shifting mosaic of growth opportunities for mobile consumers. However, a framework integrating the spatiotemporal dynamics of growth potential within riverscapes has been lacking. Here we present the concept of “foodscapes” to depict the dynamic changes in food abundance, food accessibility, and foraging costs that contribute to spatial and temporal variation of fish growth in rivers. Drawing on case studies of salmonid fishes from Alaska to California, we illustrate that “healthy” foodscapes provide a plethora of foraging opportunities—promoting diverse life history strategies that potentially enhance population stability. We identify knowledge gaps in understanding foodscapes, and approaches for management that focus on restoring trophic pathways which support diverse foraging and growth opportunities for fish in river networks.

## INTRODUCTION

To grow and survive, mobile consumers must track fluctuating resources across heterogeneous environments, sometimes over vast distances (Sinclair 2021, McNaughton et al. 1989, Abrahms et al. 2021, van der Graaf et al. 2006). While some species migrate from the north pole to the south pole (Fijn et al. 2013), and others move 1000s of km annually (Joly et al. 2019), few encounter such extreme variation in habitats as migratory salmonids. As juveniles, Pacific and Atlantic salmon, and other migratory salmonids may use all types of aquatic habitats available in large watersheds, with many species emerging in headwaters to rear in tributaries before moving into mainstem rivers or lakes. They may cross inundated floodplains and off-channel habitats, and then pass through estuaries and saltwater sloughs, to the near-marine environments and open ocean. Each of these habitats offers food to salmon critical to their growth and, ultimately, survival. But the timing (seasonal, daily, hourly, etc.) of food resource peaks—and the costs of foraging—can vary widely. How do mobile foragers like migratory salmonids make use of these dynamic patterns of food availability across the watershed habitat mosaic? What factors constrain their ability to track feeding bonanzas at the watershed scale? And finally, how can humans perceive and manage these dynamics to benefit mobile consumers like salmon, the ecosystems they inhabit, and ourselves?

The importance of physical heterogeneity across watersheds for fish has long been recognized (Schlosser 1991). Fausch et al. (2002) and Wiens (2002), adapting concepts from landscape ecology, encouraged moving beyond studies of isolated stream reaches to consider the riverscape: the continuous, hierarchical, and heterogeneous nature of riverine ecosystems. The riverscape concept, along with similar frameworks (e.g., ‘shifting habitat mosaics’ Stanford 2005) emphasize the diversity and connectivity of habitats that fish use to complete their life cycles. These frameworks have generally focused on *physical* habitat conditions, such as channel form and water temperature (but see Poff and Huryn 1998). Yet, the persistence and success of fish in these dynamic environments also depends on their ability to grow, which is a function of physical conditions, but also biotic conditions, including the abundance, quality, and accessibility of prey (Chapman 1966). It is also well-recognized that different habitats within watersheds can have distinct food sources (or ‘resource sheds’, Power and Rainey 2000) and diverse trophic pathways that support fish growth (Wipfli and Baxter 2010). For example, riverine fishes are frequently supported by a combination of instream prey resources (Bellmore et al. 2013, Rossi et. al. 2022), terrestrial invertebrates (e.g., Nakano and Murakami 2001), and marine-derived nutrients, tissue, and eggs delivered by anadromous fish (Bilby et al. 1996, Kaylor et al. 2021). Food fluxes through these pathways vary through time and may occur as pulses that, while short in duration, can be dominant sources of annual energy intake (Bently et al. 2012; Armstrong and Bond 2013).

In addition to food sources, an individual’s growth depends on food accessibility and the metabolic costs of acquiring and assimilating food (**box 1**) (Warren and Davis 1967, Fausch 1984, Hughes and Dill 1990, Piccolo et al. 2014, Naman et al. 2018 and 2019, Armstrong et al. 2021, Kaylor et al. 2021, Rossi et al. 2022). Food accessibility can change with abiotic conditions; for example, high velocities s preclude drift-foraging for small fish in larger mainstem rivers (Rosenfeld et al. 2007) and high turbidity reduces foraging efficiency of visual predators (Harvey and Railsback 2014, Piccolo et al. 2014). Some prey are not available in the water column, or have structural defenses that preclude consumption or assimilation (Wootton et al. 1996). Prey ‘accessibility’ (*sensu lato,* Railsback et al. 2005) for a given forager depends on prey morphological and behavioral traits, environmental factors, and their combined influence on tradeoffs involving risk of predation or prey handling time. Growth is also mediated by a consumer’s metabolic rate and the metabolic costs of foraging. As poikilotherms, salmon’s metabolic costs increase exponentially with temperature, which has a hump-shaped relationship with digestive capacity (Brett et al. 1969). Metabolic costs also scale with swimming exertion, affected by hydraulic conditions (e.g., velocity, turbulence) (Oldham et al. 2019). Collectively, food abundance, food accessibility, and metabolic costs all vary across space and time (**box 1**) creating a dynamic mosaic of growth hot spots that mobile consumers can exploit, albeit a mosaic in which profitability (growth potential) is offset by costs and challenges (i.e., energy expended during movement, foraging, and barriers that block movements).

Here, we synthesize these important factors and interactions into the concept of *foodscapes* – the spatial and temporal mosaic of growth potential that consumers exploit across watershed habitats (**box 2**). While a growing body of work illustrates the emerging value of a foodscape perspective in salmon ecology and management (Searle et al. 2007, Bellmore et al. 2013, Rossi 2020, Cordoleani et al. 2022, Sturrock et al. 2021, Wipfli and Baxter 2010, Armstrong et al. 2021), we are unaware of any studies that have attempted to quantify the foodscape at spatial-temporal scales that match complete salmon life histories. In effect, salmonid growth potential has not been estimated in any system at the riverscape scale (e.g., headwaters to sea) throughout the complete annual cycle. In addition, while the importance of growth potential and food for mobile consumers is well recognized, there is no holistic framework that integrates the mechanisms underlying growth potential with the concepts of resource tracking and life history diversity. Integrating concepts of growth hotspots, spatial-temporal asynchronies in growth potential, and resource tracking by salmonids with different life histories may advance our ability to apply knowledge of salmon ecology to riverscape conservation and management (See “Managing Foodscapes” below).

### Box 1.

**The spatial phenologies of three foodscape axes.**

**Figure B1.**
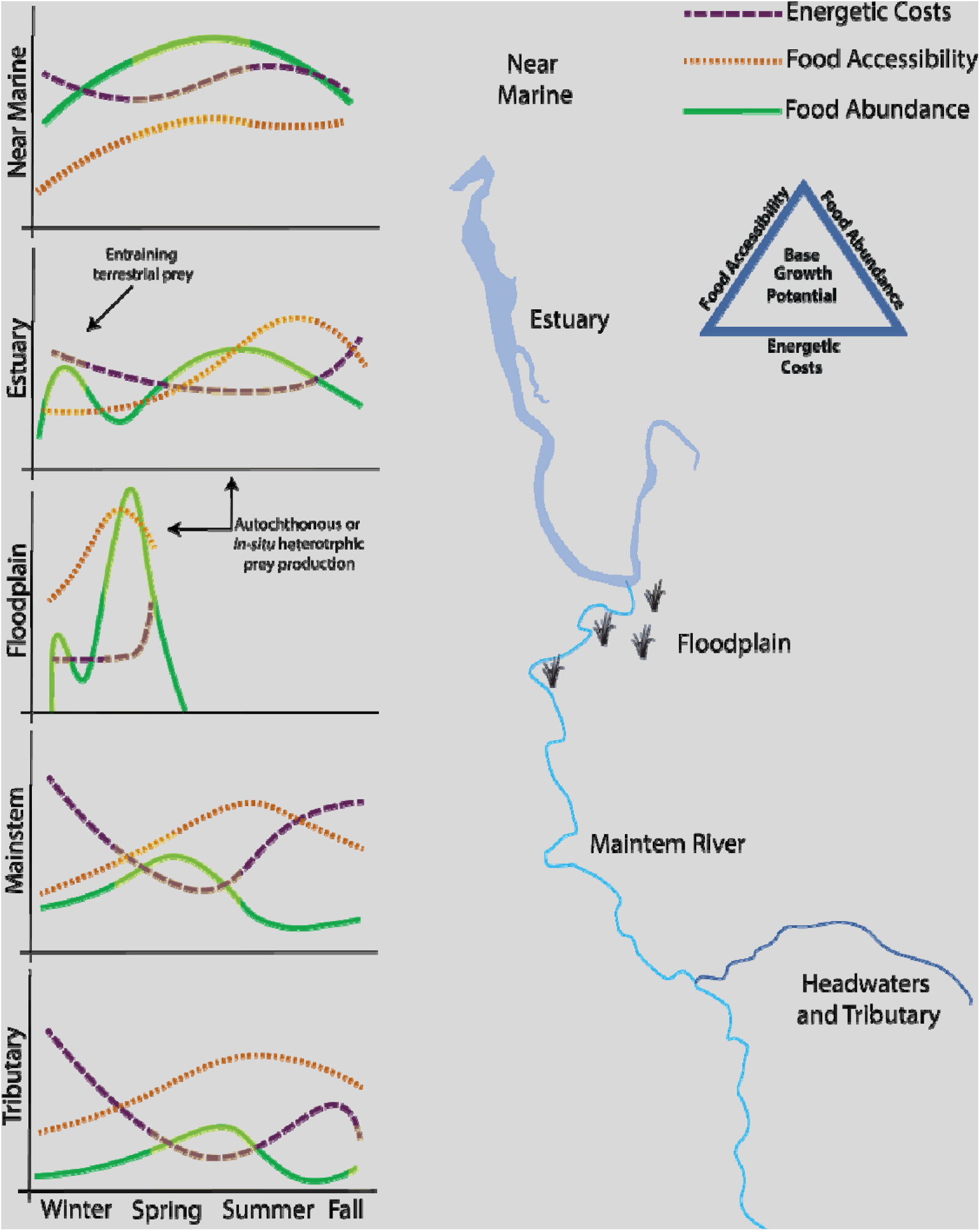
A conceptual illustration of how different seasonal patterns (phenologies) of food production, food accessibility, and energetic costs can vary across habitats of a rainfall dominated coastal riverscape leading to a seasonal-spatial mosaic of growth potential (yellow boxes show periods of high growth). We define three primary axes of the foodscape that interactively determine the base growth potential of an individual in a given habitat. Food abundance is defined as the habitat specific concentration or biomass of prey which is either produced in situ or through resource subsidies. Food accessibility is defined as the proportion of food abundance that consumers can access and is determined by both prey and habitat features (biotic and abiotic) that affect the consumer’s ability to encounter, detect, and capture a prey item with a given abundance. Energetic costs are defined as the metabolic costs of foraging, which also vary with physical (e.g., water temperature and dissolved oxygen) and biotic (e.g., predator avoidance and direct competition) factors. Endogenous factors that influence growth potential, such as aerobic scope and digestive capacity are not explicitly addressed here but can be grouped loosely under energetics. Many of these factors have been integrated in the field of foraging theory and bioenergetics (e.g. Hughes and Dill 1990) but not applied at the riverscape scale. In addition, the trophic pathways (primary to secondary production) that produce food, and the physical and biotic factors that control food accessibility and energetic costs vary among habitats and across time.

### Box 2.

**Consumer life histories interact with foodscapes to create complex growth trajectories.**

**Figure B2.**
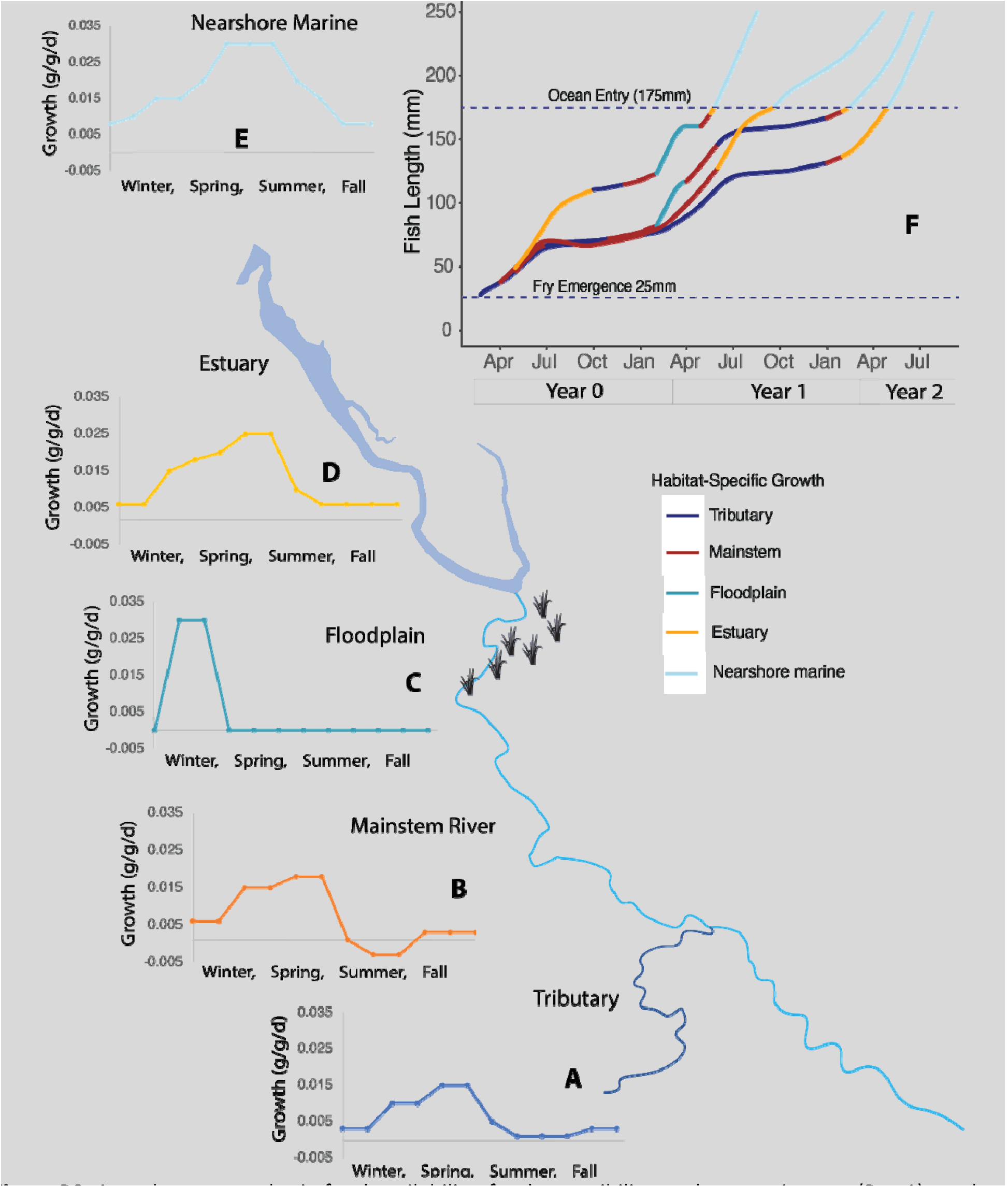
Asynchronous cycles in food availability, food accessibility, and energetic costs (Box 1) result in distinct patterns of fish growth potential across the riverscape (Panels a-e). Individual fish with different seasonal-spatial patterns of habitat use (life histories strategies) will experience different growth trajectories (panel f). Here we illustrate how four different life histories strategies for juvenile steelhead (O. mykiss) lead to different growth trajectories (panel f) using a simple bioenergetic growth model. The colors along each growth trajectory reflect the time spent in each part of the watershed from fry emergence to ocean entry. We constrained the model so that fish entered the ocean at 175 mm to illustrate how different habitat use patterns and associated growth trajectories could lead to complexity in the age and timing of ocean entry. In reality, size at ocean entry would vary, and the number of individuals that each life history supports would be a product of the habitat capacity (space) and mortality risks associated with each life history pathway which are not shown on this figure. This heuristic simulation illustrates how a complex portfolio of growth trajectories can emerge from an intact foodscape, which in turn promotes population stability in the face of environmental change (Schindler et al. 2010, Moore et al. 2014).

The growth phenologies used in this heuristic simulation were informed by measurements of seasonal and habitat-specific growth of O.mykiss in several coastal, rainfall dominated streams of the Pacific Northwest (Thompson and Beauchamp 2016, Tattam et al. 2016, Hayes et al. 2008, Stillwater 2007, and Rossi et al. 2022). However, as described, we could find no comprehensive estimates of seasonal-spatial growth patterns in any single river system– underscoring the importance of this simulation exercise. O. mykiss exhibit the most complex life histories of any Pacific salmonid, and the four life history trajectories that we used in this model were broadly taken from Shapovalov and Taft (1954), Erman & Hawthorne (1976), and Hayes et al. (2008). These included two life histories where fish reared for one years in their natal tributary, a third where fish dispersed early to an upper mainstem habitat to rear, and a fourth where young of year fish immediately moved to lower mainstem and estuarine habitat to rear (panel f).

Growth trajectories (panel f) were simulated by combining seasonal growth potentials (panels a-e) with bioenergetic modeling that accounted for allometric effects on fish metabolism (Fitzgerald et al. 2023). As fish moved between different habitats, daily mass-specific growth (g<t)) was a function of growth-rates specified for the occupied habitat i at time t (g_habitati,t_ ; panels a-e) corrected for shifting allometric constraints on growth as fish size changed through time:

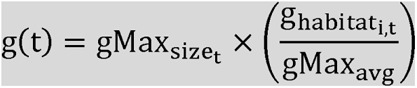

Specifically, habitat specific growth rates (g_habitati,t_) were divided by the idealized maximum growth potential of an “average” sized juvenile steelhead gMax_avg_ (assumed to be 4.5 grams), at optimum water temperature (16°C) and maximum ration (proportion Cmax = 1) using Wisconsin Bioenergetic model equations parameterized for O. mykiss (Hansen et al. 1997). The resulting quotient represents the proportion of maximum growth juvenile O. mykiss could achieve in habitat i at time t, assuming observed growth potential patterns shown in panels a-e represent those on an average sized fish. This value was then multiplied by the size-corrected maximum growth potential calculated for fish at time t (g_Maxsize__t_); a value which was recalculated each day in the bioenergetic model as fish grew. Total growth was calculated by multiplying mass-specific growth rates (g<t)) by fish mass (grams) at time t. All simulations were run on a daily time-step with a starting fish mass of 0.3 grams (approximate weight of emergent steelhead fry).

We begin by (1) defining the foodscape concept and explain how it can be applied to better understand the distribution, abundance, movement, and life history diversity of juvenile salmonids. We next (2) present case studies from Alaska to California that reveal local foodscapes and illustrate, in the aggregate, the utility of the foodscape perspective in understanding salmon population ecology. We (3) discuss approaches to evaluate foodscapes and identify knowledge gaps that currently limit our capacity to measure, model, and manage foodscapes for salmon and other mobile consumers. Finally, (4) we place foodscapes into a management context and suggest approaches for managers to incorporate a foodscape perspective into salmonid conservation and recovery planning.

## WHAT IS THE FOODSCAPE OF A MOBILE FORAGER?

We define the foodscape as a dynamic mosaic of linked habitats with different growth potential phenologies that is exploited by mobile consumers and supports multiple life histories, often through asynchronies in resource availability (**box 2**). Conceptualizing watersheds as foodscapes raises new questions about the factors that control the distribution, abundance, and life histories of mobile consumers. Some examples are: (1) How is growth potential spatially distributed across a riverscape – are there growth ‘hotspots’ and ‘hot moments’ and if so, what causes their appearance and disappearance over space and time? (2) To what extent, if any, do mobile consumers track growth potential seasonally and spatially? (3) How do the spatial-temporal patterns of growth potential support diverse life histories of consumers? And (4) which biotic interactions, and physical constraints—including those affected or effected by humans— facilitate or impair the ability of mobile consumers to exploit the foodscape? Addressing these questions can help guide strategies for managing or restoring populations that focus on supporting functional foodscapes.

From these questions we have developed three hypotheses that frame the foodscapes concept, which are presented visually in **boxes 1 and 2**.

1. *Growth potential hotspots arise in different habitats at different times due to asynchronous cycles in food availability, food accessibility, and metabolic costs*.
2. *Fish can profit by tracking peaks in growth potential across riverscapes*.
3. *Diverse foodscapes beget diverse life history strategies for mobile consumers that use the spatial compliment of habitats and food webs in different ways*.

Below we present three case studies that highlights each of these hypotheses and their importance to the foodscape concept.

## CASE STUDIES OF RIVERINE FOODSCAPES

**Hypothesis 1:** Growth potential hotspots arise in different habitats at different times due to asynchronous cycles in food availability, food accessibility, and metabolic costs: *Managed Wetlands in the Lower Sacramento River*

The Central Valley of California is a highly modified landscape with flow modification from upstream reservoirs and floodplain disconnection from a large system of levees. Few locations with river-floodplain connection remain. One of these is the Yolo Bypass, a 24,000 hectare flood bypass that routes floodwaters around the city of Sacramento. This habitat is a mosaic of agricultural fields and managed wetlands during the non-flood season. Following major precipitation events in the winter and spring months, flows overtop managed levels or weirs, generating lentic habitat with warmer temperatures and high nutrient availability that become swamped with abundant zooplankton (figure 1). Juvenile salmon exploiting these habitats grow rapidly and become much larger than individuals that remain in the mainstem (Sommer et al. 2001, Katz et al. 2017, Jeffres et al. 2020). In addition to *in situ* food production for juvenile salmon, this floodplain exports prey to downstream mainstem habitats during flood events (Sturrock et al. 2022). In turn, rapid growth and large body mass prior to ocean entry greatly enhances survival and early marine rearing of juvenile salmon (Woodson et al. 2013), suggesting that rapid growth on inundated floodplains may have population consequences. Floodplains are an example of ephemeral, yet exceptionally productive habitat that can contribute to the portfolio of successful life history strategies for juvenile salmonids.

**Figure 1.**
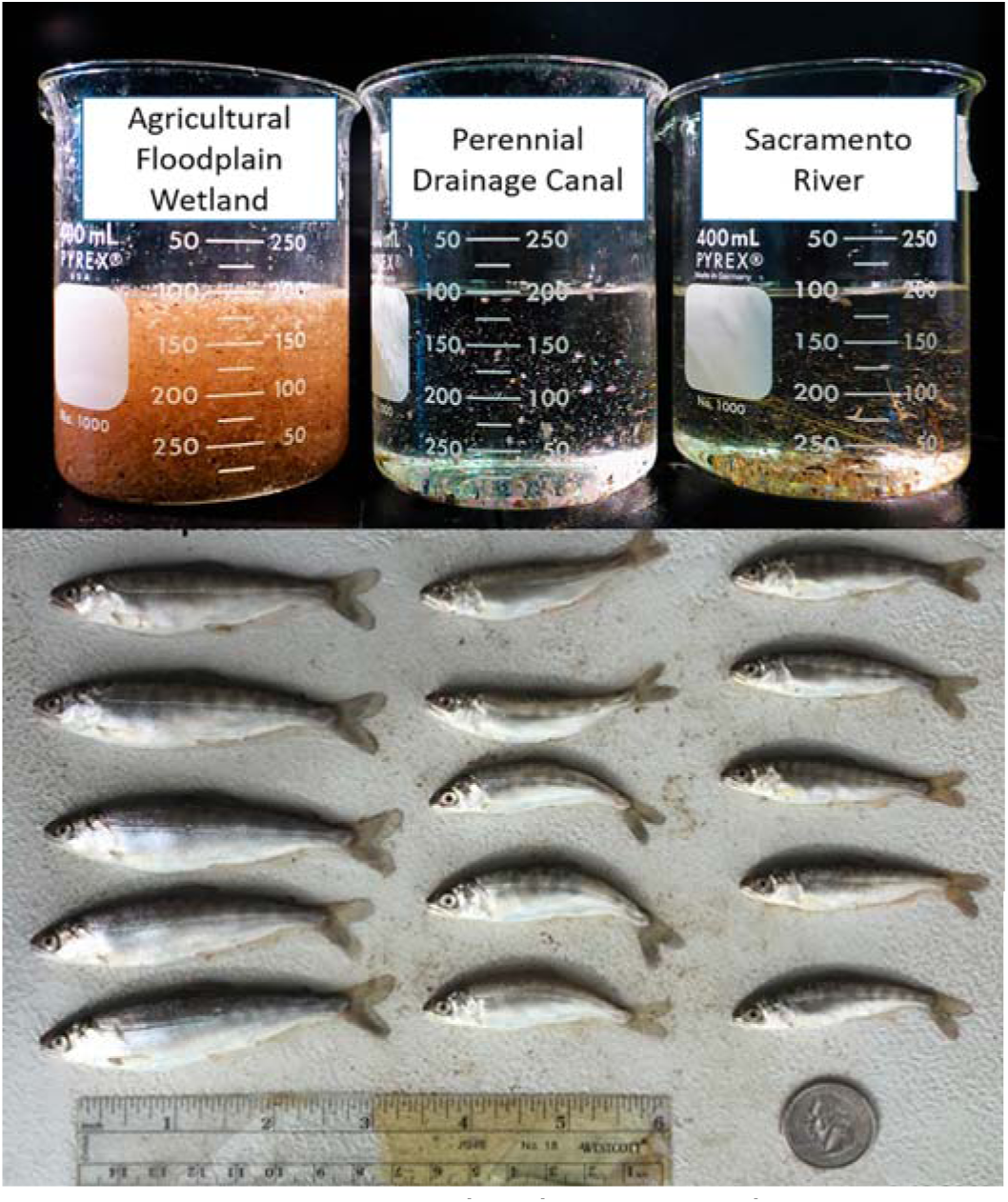
Photo of zooplankton samples and representative fish from each of the three habitat types: Flooded agricultural wetland, perennial drainage canal, and mainstem Sacramento River (figure from Jeffres et al. 2020). Note the greater abundance of zooplankton and larger associated size of fish from the flooded agricultural floodplain habitat compared to the two other locations.

The work on managed floodplains in California’s Central Valley highlights that food resource hotspots can vary dramatically across space and may be only briefly available (e.g., eggs of spawning fish, synchronous insect emergences). Many river systems have seasonal floodplains and off-channel habitats that are dry or disconnected much of the year. When inundated, these habitats provide a variety of highly productive trophic pathways for mobile consumers. As flood water fills terrestrial habitats, terrestrial invertebrates trapped by rising water can be flushed back into the main channel. Reduced floodplain velocities settle suspended sediments, increasing light penetration and stimulating primary production (Ahearn et al. 2006). In turn, primary production fuels abundant zooplankton and macroinvertebrate populations: often floodplain-specific taxa and distinct trophic pathways that support fish (Benigno and Sommer 2007, Bellmore et al. 2013, Jeffres et al. 2020, Corline et al. 2021). Although nutrient depletion may eventually diminish primary production, continued flooding can result in productive heterotrophic food web pathways supported by dissolved and particulate organic carbon liberated from inundated soils (Jeffres et al. 2020). Both primary production and heterotrophic liberation of carbon can support prolific densities of zooplankton and macroinvertebrates, an important food resource to juvenile salmonids (Sommer et al. 2001, Jeffres et al. 2008, Jeffres et al. 2020). Moreover, downstream movement of migratory fish such as juvenile salmon (from headwater habitats) frequently coincides with seasonal productivity peaks on floodplains, providing opportunities for fishes to actively or passively move into these productive off-channel habitats during periods when main-stem productivity may be low (**box 2**; Sommer et al. 2001, Jeffres et al. 2008).

**Hypothesis 2:** Fish can profit by tracking peaks in growth potential across riverscapes: *Upper Klamath Lake (Oregon) and its tributaries)*

Upper Klamath Lake is large (∼270 km^2^), shallow, and hypereutrophic. In summer, water temperatures exceed 25°C and massive cyanobacteria blooms generate poor water quality and associated fish kills. However, radio telemetry of adult redband trout revealed that these large-bodied fish migrated to the lake during fall and spring, when dissolved oxygen and pH were suitable, and temperatures were optimal for growth. In the lake, redband trout gorged on abundant chub and sculpin, feeding near their assimilative capacity. Fish moved to cool tributaries during summer and their energetic condition declined as they switched to invertebrate prey and failed to meet their metabolic demands (figure 2). After foraging in the lake during fall, fish again moved to tributaries to spawn, presumably incurring another period of energy loss.

**Figure 2.**
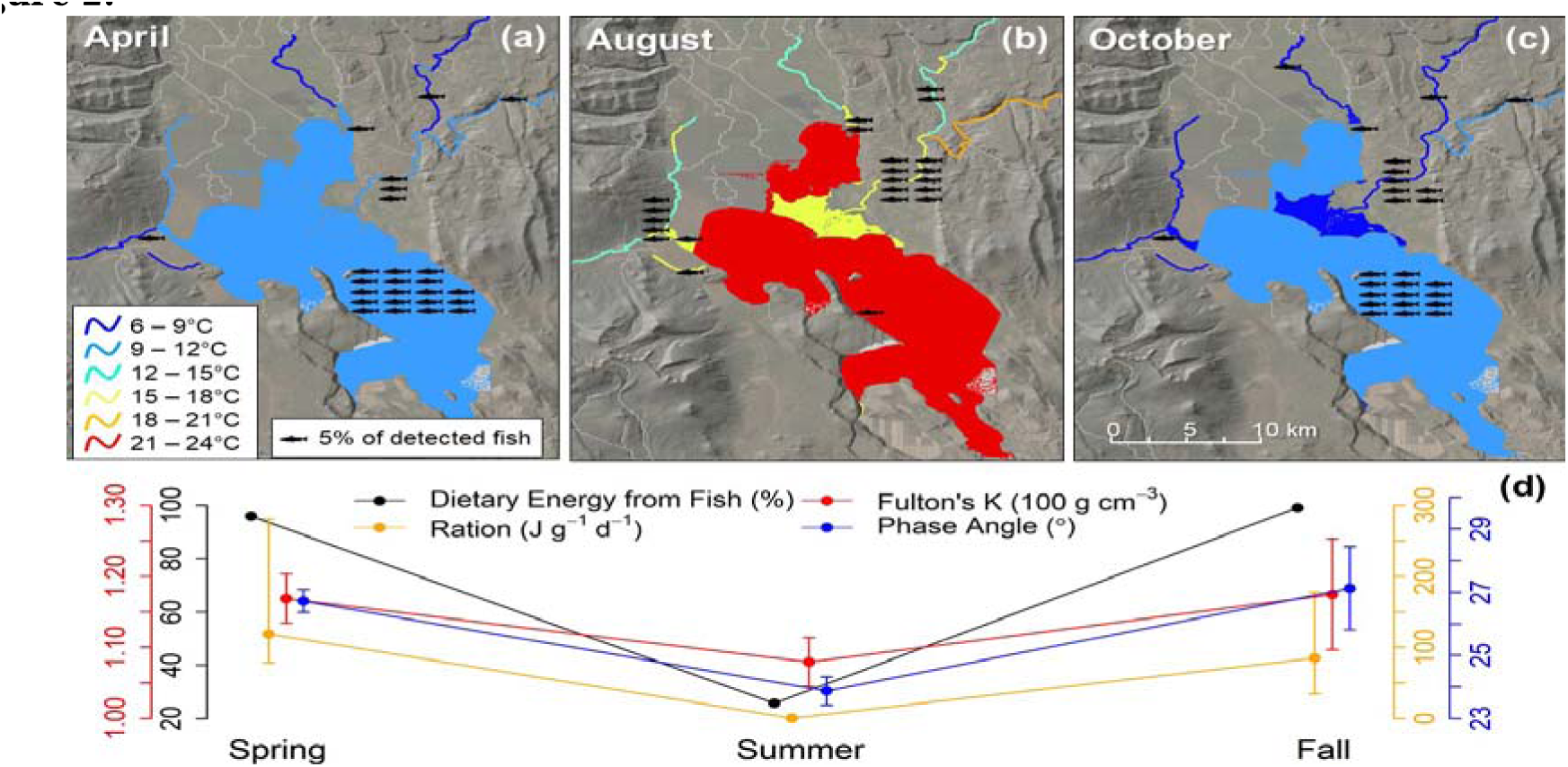
Top panel: Map of the upper Klamath Basin with monthly mean water temperatures and redband trout habitat use during April, August, and October. Habitat use sums to >100% because some fish used multiple habitats in a month. Lower panel: Patterns in foraging and energetic condition of redband trout in spring (in the lake), summer (in tributaries), and fall (in the lake). Each metric and its axis are color-coordinated (see legend). For energetic condition metrics (K and phase angle), values expressed are mean (points) and normal 95% confidence intervals(bars).” Fulton’s condition factor (K), a metric based on the ratio of weight to cubed length, and phase angle, a novel metric based on electrical properties of tissue, correlate positively and imperfectly with fish condition at different timescales. From Hahlbeck et al. (2021).

Thus, seasonally warm habitats fueled most fisheries production. This example illustrates how non-anadromous salmonids can track optimal growth conditions across large spatial scales (10s to 100s of km) and how temporal variation in water quality may mediate access to productive foraging habitat. The lake likely offers abundant food year-round, but physiological scope for growth in salmonids occurs only as pulses in spring and fall, as the lake is transitioning between suboptimally cold and warm temperatures.

Mobile consumers can track spatio-temporal variation in growth potential (Fretwell 1972, Schindler et al. 2013) with several examples of stream-dwelling fishes tracking resources across multiple habitat scales (Power 1984, Ruff et al. 2011, Armstrong & Schindler 2013). The temporal component of a habitat’s growth potential for salmonids may vary seasonally (Rossi et al. 2022) or over much shorter time frames (Armstrong et al 2013A, Baldock et al. 2015) particularly when there are trade-offs between food abundance and physiological suitability.

Salmonid behaviors like habitat cycling can alleviate these trade-offs. Feeding forays exemplify habitat cycling at timescales of minutes to hours. For example, juvenile coho salmon foray into sub-optimally cold thalweg habitat to feed on abundant salmon eggs (Armstrong et al. 2013A) or benthic invertebrates (Baldock et al. 2015) and then move to warmer floodplain or beaver-meadow complex habitat to assimilate their food. Fish similarly make forays into food-rich habitat that is overly warm (Munson et al. 1980, Sims et al. 2006) or hypoxic (Rahel and Nutzman 1994). To reduce physiological stress, individuals may exploit diel variation and enter foraging habitats during more favorable periods (Sims et al. 2006), or leave before physiological costs accrue (Rahel and Nutzman 1994, Pepino et al. 2015). Similarly, diel vertical movements of juvenile sockeye salmon track intermediate light levels (the anti-predation window) and fish adjust their cyclical movements as day lengths and light penetration vary intra-seasonally (Armstrong et al 2013B). At coarser timescales, salmonids may move seasonally between refuge and foraging habitats. For example, lake trout are coldwater specialists restricted to deeper pelagic habitat during summer (Martin 1952, Guzzo 2017). However, in spring and fall, when lake conditions are relatively isothermal and cool, these fish exploit productive littoral habitats (Martin 1952, Guzzo et al. 2017). Indeed, water bodies that would be lethal to salmonids in summer may fuel a substantial fraction of fish production, as demonstrated by recent work (figure 2, Hahlbeck et al. 2021).

**Hypothesis 3:** Diverse foodscapes beget diverse life history strategies for mobile consumers that use the spatial complement of habitats and food webs in different ways: *A coastal watershed in the Gulf of Alaska*

In the Gulf of Alaska, coastal watersheds frequently contain a mosaic of glacier-, snow-, and rain-fed tributaries (figure 3a) that have distinct, and frequently asynchronous seasonal flow, temperature, and nutrient regimes (Hood and Berner 2009, Fellman et al. 2014). These pronounced physicochemical differences drive divergent seasonal cycles of production and availability of aquatic resources that support juvenile salmon (Bellmore et al. 2022), creating foraging and growth “hotspots” at different times of the year (figure 3b). Resource availability was predicted to be highest in glacier-fed streams in the spring, rain-fed streams in the summer, and snow-fed streams in the autumn. Bellmore et al. (2022) showed that juvenile salmon to move among glacier-, snow-, and rain-fed streams, tracking these seasonal asynchronies in food availability and energetically favorable foraging and growth opportunities as they shift through time across the river network (figure 3c,d). These behavioral options led to a diversity of different growth trajectories (Bellmore et al. 2022) that likely contribute to population resilience; i.e., different habitat use patterns that succeed in one year may not be favored in another. As glaciers and snow disappear from the landscape, however, the hydrology of watersheds will become more homogeneous (Barnett et al. 2005, O’Neel et al. 2015). This hydrologic homogenization could shrink foraging and growth opportunities as seasonal patterns of resource availability across the foodscape become more synchronous, narrowing the range of habitat use patterns and associated growth trajectories that juvenile salmon can express (figure 3d).

**Figure 3.**
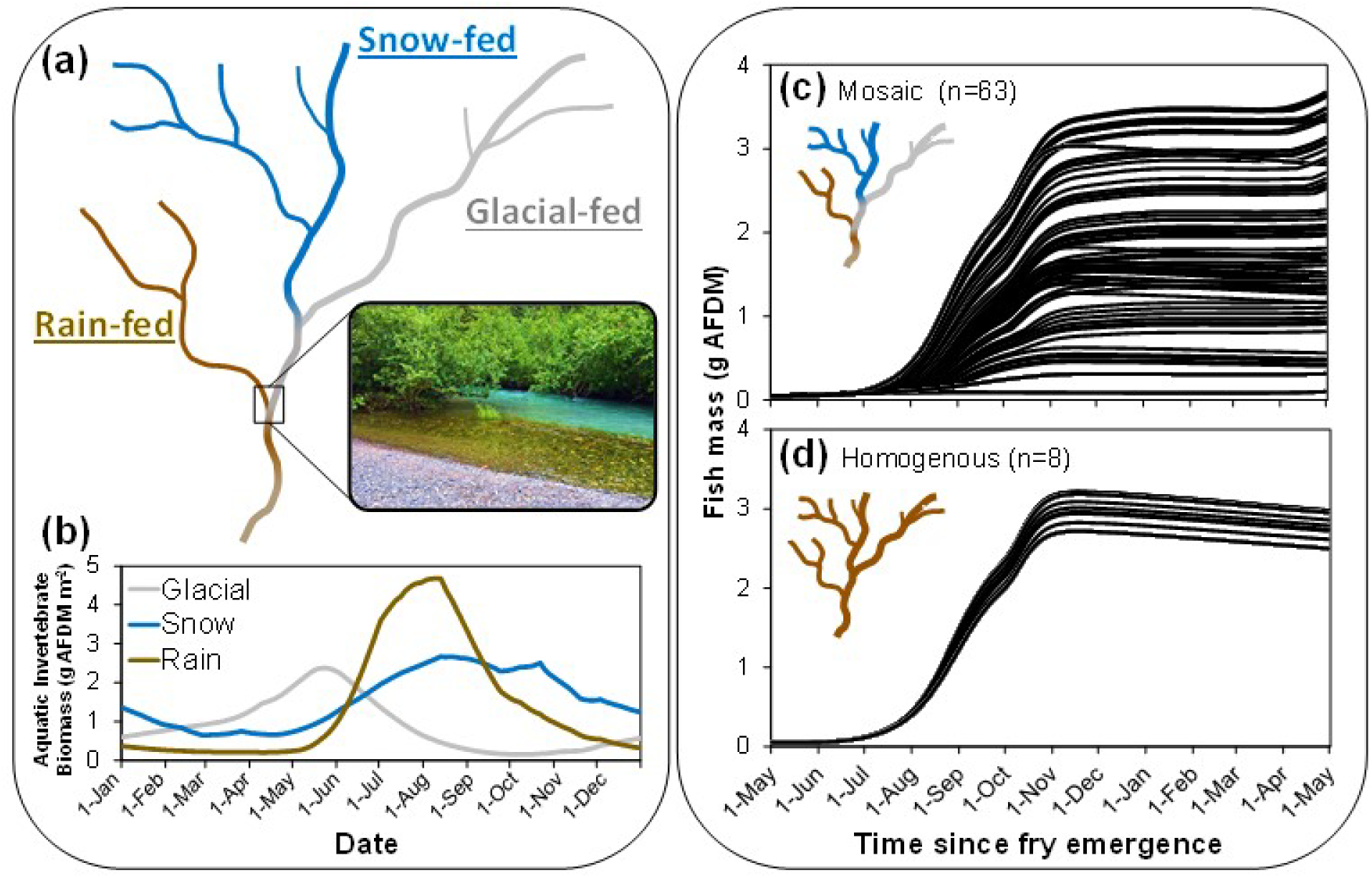
Modified from Bellmore et al. (2022), mosaics of glacier-, snow-, and rain-fed streams in Coastal Gulf of Alaska watersheds create diverse foodscapes for juvenile salmon. Panel A is a conceptual river network diagram with an inset picture that shows the confluence of a tannin-stained rain-fed streams with cold and turbid glacier-fed streams. Panel B: modeled patterns of aquatic invertebrate prey availability supported in each stream type. Panels C and D: juvenile salmon growth trajectories supported in a diverse river network (“mosaic”) that contains all three stream types, and a homogenous network composed entirely of rain-fed streams. Simulations conducted with an individual based model initialized with 200 juvenile fish with different movement propensities. AFDM = ash-free-dry-mass.

Life history diversity has been key to the resilience of Pacific salmon (Schindler et al. 2010; Carlson & Satterthwaite 2011). Pacific salmon life-history diversity is most frequently associated with variation in the age, timing, and size of transitions between life-stages (i.e., smolting and maturation; Quinn, 2005), which has been shown to stabilize population dynamics via “portfolio effects” (Moore et al., 2014, Schindler et al., 2010). But here we define life history as also including variation in movement and associated habitat use patterns that salmon and trout have been shown to exhibit during their freshwater life stages (**box 2**; Koski 2009, Bourret et al. 2016),. These can influence their growth, survival, and subsequent reproductive success. Variation within a watershed in the availability of mainstem, tributary, floodplain, and estuary habitats provide numerous permutations of habitat use, foraging opportunities, and growth potentials in foodscapes, differentiating growth trajectories in ways that may promote life history diversity and population resilience (see **box 2**). As this case study illustrates, hydrologic heterogeneity within watersheds, such as tributaries with different runoff patterns, may also promote life history diversity (see also Dralle et al. 2023).

## KNOWLEDGE GAPS IN FOODSCAPES

Certain aspects of the foodscape can be measured and modeled using established methods. For example, process-based and statistical approaches used to estimate water temperature across whole watersheds are rapidly expanding (Isaak et al. 2017, Fullerton et al. 2015). Similarly, methods to estimate channel hydraulics and light inputs across wide scales are becoming increasingly accessible (Tamminga et al. 2015, Savoy et al. 2019, Bode et al. 2010), as are bio-logging tags that can quantify metabolic costs (e.g. Lennox et al. 2019). Promising ecological modeling approaches that can estimate food availability and growth potential from known channel and riparian conditions are also emerging (Bellmore et al. 2017), which could be expanded to networks scales (e.g., Bellmore et al. 2022). Other aspects of foodscapes are less well understood and difficult to estimate. Below we outline key knowledge gaps that, if filled, would help us better understand and quantify foodscapes.

### Watershed-scale estimates of food availability and quality

Prey availability is often measured at small spatial (e.g., reach) and narrow temporal (e.g. seasonal, or single sampling event) scales, limiting our understanding of prey abundance across larger scales of time and space. Due to the effort required for macroinvertebrate sampling and processing, studies focused on capturing spatial patterns of prey availability often do so at the cost of temporal replication, and vice versa. Few studies have attempted to quantify prey availability across a range of available habitats and over time (but see Bellmore et al. 2013 and Cordoleani et al. 2022). In addition, information on food availability and production is often descriptive, so we lack predictive tools to estimate prey as we estimate physical parameters such as temperature (but see Bellmore et al. 2017). What would a watershed-scale model of food production and quality look like?

While estimating prey availability precisely at fine scales is highly complex (e.g., Hayes et al. 2018) and may not be tractable in many situations, coarser-scale heuristic models could likely capture broad differences among habitats (Bellmore et al. 2017, Whitney et al. 2020).

Alternatively, proxies of food production could be employed. Estimates of primary production derived from open channel metabolism measurements (Hall and Hotchkiss 2017) could be used to predict the amounts of basal resources fixed in different habitats, seasonal variation in biological productivity (Mejia et al. 2019), and the energetic capacity to support fishes (McGarvey and Johnston 2011). While this approach may not be applicable at fine spatial scales, it could be useful for predicting broad patterns in biological productivity across watershed networks (Ruegg et al. 2021).

In addition to prey abundance, there is also proliferating evidence that food quality (nutritional content) is critical for growth and development of consumers (Twinning et al. 2016). Emerging techniques using fatty acid profiles to estimate prey quality suggest that differences in prey quality, especially between aquatic and terrestrial prey sources, can impose important nutritional bottlenecks on fish (Zavorka et al. 2021). However, we lack a broad understanding of how prey quality varies at watershed scales through time. Furthermore, the quality of prey (independent of quantity) may influence foraging and movement decisions by consumers in ways that aren’t fully understood (Twining et al. 2021); e.g., fishes could make directed movements to habitats where prey abundance is low but nutritional quality is high. Thus, prey quality is an essential part of foodscapes that warrants further attention.

### Alternate foraging modes, habitats, and timescales

Work on fish foraging has not yet explored the full range of conditions fish encounter in the foodscape. First, most focus has been on drift-foraging by salmonids (e.g., Hughes and Dill 1990, Piccolo et al. 2014). While this is the dominant strategy in flowing water, salmonids frequently forage in other ways, especially in more lentic habitats such as floodplains (Jeffres et al. 2020) or low velocity pools (Harvey and Railsback 2014, Naman et al. 2018, Rossi et al. 2021). Factors that constrain prey accessibility and energetic costs in these habitats are less understood and would benefit from a similar level of attention that has been given to drift-foraging. Second, foraging observations have been almost entirely constrained to smaller habitats accessible to humans. Foraging in larger mainstem rivers is largely unknown, despite well-documented occurrences of fish in these habitats (e.g., Bradford and Taylor 1997). How fish ‘make a living’ in these bigger systems in the face of harsh hydraulic conditions, often high turbidity, warm summer temperatures, and higher predation risk is a key gap in our understanding of the foodscape. Third, studies characterizing foraging behavior in streams have been limited to summer and low flows (but see Nakano 1999, Nielsen 1990), which does not represent the full range of conditions they experience. High flow conditions may provide important foraging opportunities for some species by expanding inundated environments and entraining terrestrial and aquatic prey into the water column (Fitzgerald et al. 2023). Finally, fish foraging behaviors are highly context dependent, influenced by food availability but also by a suite of other factors that vary across the riverscape and over time, including physical habitat quality, predation risk, and inter- and intraspecific competition (White et al. 2014). Expanding empirical studies and theory to include more of the foraging conditions fish actually experience in watersheds is critical to better understanding the foodscape.

### Resource tracking by fish

A central tenet of the foodscape concept is that fish can, and often do, track spatio-temporal variation in growth potential. While there is evidence to support this (e.g., Ruff et al. 2011, Armstrong et al. 2013A, Hansen and Closs 2009, Power 1984), the mechanisms underlying resource tracking are less understood. What cues do fish use to track resources? Over what spatiotemporal scales does this occur? And what are the population and food web consequences of resource tracking? While broader ecological theory has advanced mechanistic understanding of these processes (e.g., Mueller and Fagan 2008, Fagan et al. 2017), applications to aquatic ecosystems are limited (Johnson and Rice 2014).

Fish have neither omniscient knowledge of their environments nor perfect access to the best available at any time, so resource tracking is imperfect. Understanding when and why resource tracking fails (and the consequences on consumers) is critical for understanding foodscape dynamics. A particularly important question is how anthropogenic alterations to watersheds affect resource tracking. For example, physical habitat alteration could disrupt important cues fish use to acquire and process information about their environments, creating ecological traps where low-quality habitats are perceived as preferred and high-quality habitats are avoided (Patten and Kelly 2010, Hale and Swearer 2017). Human impacts to the fish themselves could also affect resource tracking. For instance, human-induced selection through increasing hatchery supplementation could misalign behavioral traits or spawning distributions or timing (Hoffnagle et al. 2008) with growth hotspots. This hatchery effect is poorly understood, but could severely reduce the ability of fish to exploit shifting growth opportunities, given rapid declines in wild populations (Price et al. 2020).

Resource tracking by mobile consumers may also contribute to foodscape ‘stability’. Mobile consumers “crop” resource peaks that could destabilize local food webs, and move nutrients and organic matter across watershed networks. Ecological theory suggests that these meta-food web linkages could benefit both mobile and sessile organisms by stabilizing food web dynamics (McCann et al. 2005). Conversely, reducing the extent of resource tracking could lead to destabilizing effects, e.g. via resource ‘escape’ (Ryser et al. 2021). Further empirical study is needed to test these theoretical predictions.

## MANAGING FOODSCAPES

Most salmon bearing watersheds have been degraded by humans (Lichatowich et al. 1999). A foodscape perspective to salmon recovery expands our view of restoration to encompass the sources, phenology, and flow paths of key food resources (Bellmore et al. 2017). Considering foodscapes in management also focuses our attention on maintaining conditions that allow populations of salmonids to track and exploit optimal feeding opportunities across the riverscape (White et al. 2014). Frequently, restoration objectives for salmon species are based on population targets and physical habitat suitability metrics, but these approaches have been criticized for failing to protect vital ecological processes and favoring heavily managed solutions that can result in unnaturally static conditions (Trush et al. 2000, Anderson et al. 2006, Beechie et al. 2010). In recent years, process-based restoration has emerged as an alternative paradigm that addresses these concerns. Process-based restoration aims to “reestablish normative rates and magnitudes of physical and ecological processes that sustain river and floodplain ecosystems” (Beechie et al. 2010, Trush et al. 2000). However, in practice, process-based restoration has primarily emphasized reestablishment of physical (typically channel and riparian) processes, assuming—with often little evidence—that food web and consumer recovery will necessarily follow (Whitney et al. 2020). A more explicit focus on the spatio-temporal dynamics of food webs that sustain salmon (e.g. foodscapes), would benefit the field of process-based restoration.

In contrast to traditional population targets, or static form-based habitat-based restoration programs, foodscape management can be framed within process-based restoration by explicitly emphasizing the processes and interactions that control the growth of consumers across the riverscape and through time. For any given combination of (a) river system and (b) consumer, the foodscape managers can ask: how have patterns and processes affecting each of the three axes of the foodscape (food abundance, food accessibility, and the energetic costs of foraging identified in Box 1) been altered by human modification of the river system and how can they be recovered? For instance, were certain trophic pathways (from basal primary production to ingestion of prey by the consumer) lost, which, if restored, could help re-establish certain life histories of target fishes (e.g. floodplain rearing for Chinook salmon)? How should novel foodscapes, which leverage anthropogenically modified landscapes to take advantage of foraging opportunities (e.g. Cordoleani et al. 2022, Sturrock et al. 2021, Jeffres et al., 2020), be balanced against work to restore an historic foodscape? How can we compare and prioritize (or at least sequence) investment in recovering specific elements of each axis of the foodscape to improve restoration outcomes?

We acknowledge that these foodscape questions are challenging, if not impossible, to answer definitively– particularly in highly modified river systems. Quantifying *current* patterns of food production, food availability, energetic costs, and salmon life histories at a riverscape scale requires intensive sampling effort, and a suite of food web sampling methods and modeling (Naman et al. 2022), some of which have yet to be developed (see “Knowledge Gaps in Foodscapes”). Estimating how patterns of growth potential across a foodscape have been altered by human modification may require a combination of all these methods along with estimates of unimpaired physicochemical and biotic conditions, and historical and Indigenous knowledge (Quaempts et al. 2018, Atlas et al. 2021). However, we suggest that that baseline foodscape patterns *can* be reasonably estimated (at least for well-studied taxa like Pacific salmon) using a conceptual model framework – and then quantified and tested using a nested, iterative approach of modeling, monitoring, and experimentation (Power et al. 1998). Conceptual foodscape models should link a suite of predicted, even if not currently present, salmon life histories with the predicted phenologies and magnitudes of growth potential presented to them at the riverscape scale (e.g., Box 2). Conceptual foodscape models can be developed based on a combination river food web ecology, existing knowledge about common life histories of the focal prey and consumer species and how they interact with hydrology and stream type. These models help focus subsequent field work to resolve critical uncertainties, and expand inferences to broader ranges of fluctuating climatic, hydrologic, and human-imposed conditions.

Once these conceptual models are developed for both baseline and impaired foodscapes, their underlying assumptions can be tested and refined based on carefully designed monitoring studies that link growth and movement of salmonids to the spatial and temporal pattern of food production and foraging costs in a watershed (e.g. Cordoleani et al. 2022, Sturrock et al. 2021, Hahlbeck et al. 2021), with these data streams informing food web modeling efforts (e.g.

Bellmore et al. 2022). By comparing the baseline foodscape against the impaired or realized foodscape, we can identify and prioritize potential mismatches, and investigate the causes and potential solutions for those mismatches. Even with imperfect foodscape models, these questions may reveal new insights and testable hypothesis about what is limiting the capacity and diversity of salmon life histories and, thus, population resilience.

## CONCLUSION

The foodscape is a mosaic of linked habitats with different growth potential phenologies that are exploited by mobile consumers. While the constituent themes of the foodscape concept are well established (e.g., riverscapes, food webs, resource tracking), this is among the first efforts to synthesize them, particularly at spatial scales that match the life history of mobile consumers (but see Oullet et al., *in prep)* for emerging approaches to evaluating foodscapes). Here we describe the foodscape using salmonids as a focal organism, but the concept is generalizable to any mobile consumer that has the ability to track fluctuating patterns of growth potential across time and space (e.g. Sinclair 2021). Foodscapes also offer a holistic framework to integrate food web considerations into watershed management and salmon recovery, an increasingly recognized knowledge gap. The foodscape framework will generate testable hypotheses, and ultimately management actions to guide the recovery and stewardship of watersheds and freshwater organisms.

## ACKNOWLEDGMENTS

The authors would like to thank the Food For Fish working group, led by Valerie Ouellet and Aimee Fullerton (NOAA) who are developing a paper on the challenges and opportunities for quantifying foodscapes in river networks and who provided valuable feedback on this manuscript. Gabriel Rossi, Mary Power, and Stephanie Carlson were supported by a National Science Foundation CZP EAR-1331940 for the Eel River Critical Zone Observatory, and Gabriel Rossi was also supported by generous contributions from California Trout.

## DATA AVAILABILITY STATEMENT

The model and data used for Box 2 is available at: exchange.iseesystems.com/public/ryan-bellmore/fish-foodscape-ibm-example/index.html

## REFERENCES

Abrahms, B., E. O. Aikens, J. B. Armstrong, W. W. Deacy, M. J. Kauffman, and J. A. Merkle. 2021. Emerging Perspectives on Resource Tracking and Animal Movement Ecology. Trends in Ecology & Evolution 36:308–320.

Armstrong, J. B., and D. E. Schindler. 2013 A. Going with the Flow: Spatial Distributions of Juvenile Coho Salmon Track an Annually Shifting Mosaic of Water Temperature. Ecosystems 16:1429–1441.

Armstrong, J.B., Schindler, D.E., Ruff, C.P., Brooks, G.T., Bentley, K.E., Torgersen, C.E. 2013 B. Diel horizontal migration in streams: Juvenile fish exploit spatial heterogeneityin thermal and trophic resources. Ecology, 94, 2066–2075.

Armstrong, J. B., D. E. Schindler, C. P. Ruff, G. T. Brooks, K. E. Bentley, and C. E. Torgersen. 2013. Diel horizontal migration in streams: Juvenile fish exploit spatial heterogeneity in thermal and trophic resources. Ecology 94: 2066–2075.

Atlas, W. I., N. C. Ban, J. W. Moore, A. M. Tuohy, S. Greening, A. J. Reid, N. Morven, E. White, W. G. Housty, J. A. Housty, C. N. Service, L. Greba, S. Harrison, C. Sharpe, K. I. R. Butts, W. M. Shepert, E. Sweeney-Bergen, D. Macintyre, M. R. Sloat, and K. Connors. 2021. Indigenous Systems of Management for Culturally and Ecologically Resilient Pacific Salmon (*Oncorhynchus* spp.) Fisheries. BioScience 71:186–204.

Baldock, J. R., J. B. Armstrong, D. E. Schindler, and J. L. Carter. 2016. Juvenile coho salmon track a seasonally shifting thermal mosaic across a river floodplain. Freshwater Biology 61:1454–1465.

Baxter, C. V., K. D. Fausch, and W. Carl Saunders. 2005. Tangled webs: reciprocal flows of invertebrate prey link streams and riparian zones: Prey subsidies link stream and riparian food webs. Freshwater Biology 50:201–220.

Beechie, T. J., D. A. Sear, J. D. Olden, G. R. Pess, J. M. Buffington, H. Moir, P. Roni, and M. M. Pollock. 2010. Process-based Principles for Restoring River Ecosystems. BioScience 60:209–222.

Bellmore, J. R., C. V. Baxter, K. Martens, and P. J. Connolly. 2013. The floodplain food web mosaic: a study of its importance to salmon and steelhead with implications for their recovery. Ecological Applications 23:189–207.

Bellmore, J. R., J. R. Benjamin, M. Newsom, J. A. Bountry, and D. Dombroski. 2017. Incorporating food web dynamics into ecological restoration: a modeling approach for river ecosystems. Ecological Applications 27:814–832.

Bellmore, J. R., J. B. Fellman, E. Hood, M. R. Dunkle, and R. T. Edwards. 2022. A melting cryosphere constrains fish growth by synchronizing the seasonal phenology of river food webs. Global Change Biology 28:4807–4818.

Benigno, G. M., and T. R. Sommer. 2008. Just add water: sources of chironomid drift in a large river floodplain. Hydrobiologia 600:297–305.

Bentley, K. T., D. E. Schindler, J. B. Armstrong, R. Zhang, C. P. Ruff, and P. J. Lisi. 2012. Foraging and growth responses of stream-dwelling fishes to inter-annual variation in a pulsed resource subsidy. Ecosphere 3(12):1–17

Bilby, R. E., B. R. Fransen, and P. A. Bisson. 1996. Incorporation of nitrogen and carbon from spawning coho salmon into the trophic system of small streams: evidence from stable isotopes. Canadian Journal of Fisheries and Aquatic Sciences 53:164–173.

Bourret, S. L., C. C. Caudill, and M. L. Keefer. 2016. Diversity of juvenile Chinook salmon life history pathways. Reviews in Fish Biology and Fisheries 26:375–403.

Bradford, M. J., and G. C. Taylor. 1997. Individual variation in dispersal behaviour of newly emerged chinook salmon (Oncorhynchus tshawytscha) from the Upper Fraser River, British Columbia. Canadian Journal of Fisheries and Aquatic Sciences 54:1585–1592.

Brett, J. R., J. E. Shelbourn, and C. T. Shoop. 1969. Growth Rate and Body Composition of Fingerling Sockeye Salmon, *Oncorhynchus nerka*, in relation to Temperature and Ration Size. Journal of the Fisheries Research Board of Canada 26:2363–2394.

Cederholm, C. J., M. D. Kunze, T. Murota, and A. Sibatani. 1999. Pacific Salmon Carcasses: Essential Contributions of Nutrients and Energy for Aquatic and Terrestrial Ecosystems. Fisheries 24:6–15.

Cordoleani, F., Holmes E., Bell-Tilcock M., Johnson R.C., Jefres C. 2022. Variability in foodscapes and fsh growth across a habitat mosaic: implications for management and ecosystem restoration. Ecological Indicators 136:108681

Corline N.J., Peek .R.A., Montgomery J., Katz J.V.E., Jefres C.A. 2021. Understanding community assembly rules in managed foodplain food webs. Ecosphere 12(2):e03330

Dralle, D. N., G. Rossi, P. Georgakakos, W. J. Hahm, D. M. Rempe, M. Blanchard, M. E. Power, W. E. Dietrich, and S. M. Carlson. 2023. The salmonid and the subsurface: Hillslope storage capacity determines the quality and distribution of fish habitat. Ecosphere 14(2).

Fagan, W. F., E. Gurarie, S. Bewick, A. Howard, R. S. Cantrell, and C. Cosner. 2017. Perceptual Ranges, Information Gathering, and Foraging Success in Dynamic Landscapes. The American Naturalist 189:474–489.

Fausch, K. D. 1984. Profitable Stream Positions for Salmonids: Relating Specific Growth Rate to Net Energy Gain. Canadian Journal of Zoology 62: 441–51.

Fausch, K. D., C. E. Torgersen, C. V. Baxter, and H. W. Li. 2002. Landscapes to Riverscapes: Bridging the Gap between Research and Conservation of Stream Fishes. BioScience 52:483.

Fellman, J. B., E. Hood, R. G. M. Spencer, A. Stubbins, and P. A. Raymond. 2014. Watershed Glacier Coverage Influences Dissolved Organic Matter Biogeochemistry in Coastal Watersheds of Southeast Alaska. Ecosystems 17:1014–1025.

Fijn, R. C., D. Hiemstra, R. A. Phillips, and J. van der Winden. 2013. Arctic Terns *Sterna paradisaea* from the Netherlands Migrate Record Distances Across Three Oceans to Wilkes Land, East Antarctica. Ardea 101:3–12.

Fitzgerald, K. A., J. R. Bellmore, J. B. Fellman, M. L. H. Cheng, C. E. Delbecq, and J. A. Falke. 2023. *In Press*. Stream hydrology and a pulse subsidy shape patterns of fish foraging. Journal of Animal Ecology.

Fretwell, S. D., and H. L. Lucas, JR. 1970. On territorial behavior and other factors influencing habitat distribution in birds. Acta Biotheoretica 19:16–32.

Grosholz, E., and E. Gallo. 2006. The influence of flood cycle and fish predation on invertebrate production on a restored California floodplain. Hydrobiologia 568:91–109.

Guzzo, M. M., P. J. Blanchfield, and M. D. Rennie. 2017. Behavioral responses to annual temperature variation alter the dominant energy pathway, growth, and condition of a cold-water predator. Proceedings of the National Academy of Sciences 114:9912–9917.

Harvey, B. C., and S. F. Railsback. 2014. Feeding Modes in Stream Salmonid Population Models: Is Drift Feeding the Whole Story? Environmental Biology of Fishes 97: 615–25.

‘Hahlbeck, N., W. R. Tinniswood, M. R. Sloat, J. D. Ortega, M. A. Wyatt, M. E. Hereford, B. S. Ramirez, D. A. Crook, K. J. AnlaufJDunn, and J. B. Armstrong. 2022. Contribution of warm habitat to coldJwater fisheries. Conservation Biology 36.

Hale, R., and S. E. Swearer. 2017. When good animals love bad restored habitats: how maladaptive habitat selection can constrain restoration. Journal of Applied Ecology 54:1478–1486.

Hall, R. O., and E. R. Hotchkiss. 2017. Stream Metabolism. Pages 219–233 Methods in Stream Ecology. Elsevier.

Hansen, E. A., and G. P. Closs. 2009. Long-term growth and movement in relation to food supply and social status in a stream fish. Behavioral Ecology 20:616–623.

Heim, K. C., T. E. McMahon, L. Calle, M. S. Wipfli, and J. A. Falke. 2019. A general model of temporary aquatic habitat use: Water phenology as a life history filter. Fish and Fisheries:faf.12386.

Hoffnagle, T. L., R. W. Carmichael, K. A. Frenyea, and P. J. Keniry. 2008. Run Timing, Spawn Timing, and Spawning Distribution of Hatchery- and Natural-Origin Spring Chinook Salmon in the Imnaha River, Oregon. North American Journal of Fisheries Management 28:148–164.

Hood, E., and L. Berner. 2009. Effects of changing glacial coverage on the physical and biogeochemical properties of coastal streams in southeastern Alaska. Journal of Geophysical Research 114:G03001.

Hughes, N. F., and L. M. Dill. 1990. Position Choice by Drift-Feeding Salmonids: Model and Test for Arctic Grayling (*Thymallus arcticus*) in Subarctic Mountain Streams, Interior Alaska. Canadian Journal of Fisheries and Aquatic Sciences 47:2039–2048.

Jefres C.A., Opperman J.J., Moyle P.B. 2008. Ephemeral foodplain habitats provide best growth conditions for juvenile Chinook salmon in a California river. Environmental Biology of Fishes 83(4):449–458

Jeffres, C. A., E. J. Holmes, T. R. Sommer, and J. V. E. Katz. 2020. Detrital food web contributes to aquatic ecosystem productivity and rapid salmon growth in a managed floodplain. PLOS ONE 15:e0216019.

Johnson, M. F., and S. P. Rice. 2014. Animal perception in gravel-bed rivers: scales of sensing and environmental controls on sensory information. Canadian Journal of Fisheries and Aquatic Sciences 71:945–957.

Joly, K., E. Gurarie, M. S. Sorum, P. Kaczensky, M. D. Cameron, A. F. Jakes, B. L. Borg, D. Nandintsetseg, J. G. C. Hopcraft, B. Buuveibaatar, P. F. Jones, T. Mueller, C. Walzer, K. A. Olson, J. C. Payne, A. Yadamsuren, and M. Hebblewhite. 2019. Longest terrestrial migrations and movements around the world. Scientific Reports 9:15333.

Katz, J. V. E., C. Jeffres, J. L. Conrad, T. R. Sommer, J. Martinez, S. Brumbaugh, N. Corline, and P. B. Moyle. 2017. Floodplain farm fields provide novel rearing habitat for Chinook salmon. PLOS ONE 12:e0177409.

Kaylor, M. J., J. B. Armstrong, J. T. Lemanski, C. Justice, and S. M. White. 2022. Riverscape heterogeneity in estimated Chinook Salmon emergence phenology and implications for size and growth. Ecosphere 13 (7) e4160.

Koski, K. V. 2009. The Fate of Coho Salmon Nomads: The Story of an Estuarine-Rearing Strategy Promoting Resilience. Ecology and Society 14:art4.

Lennox, R. J., J. M. Chapman, W. M. Twardek, F. Broell, K. Bøe, F. G. Whoriskey, I. A. Fleming, M. Robertson, and S. J. Cooke. 2019. Biologging in combination with biotelemetry reveals behavior of Atlantic salmon following exposure to capture and handling stressors. Canadian Journal of Fisheries and Aquatic Sciences 76:2176–2183.

Lichatowich, J., Mobrand, L., and Lestelle, L. 1999. Depletion and extinction of Pacific salmon (Oncorhynchus spp.): A different perspective. – ICES Journal of Marine Science, 56: 467–472

Martin, N. V. 1952. A Study of the Lake Trout, Salvelinus Namaycush, in two Algonquin Park, Ontario, Lakes. Transactions of the American Fisheries Society 81:111–137.

McCann, K. S., J. B. Rasmussen, and J. Umbanhowar. 2005. The dynamics of spatially coupled food webs: Spatially coupled food webs. Ecology Letters 8:513–523.

McGarvey, D. J., and J. M. Johnston. 2011. A Simple Method to Predict Regional Fish Abundance: An Example in the McKenzie River Basin, Oregon. Fisheries 36:534–546.

McNaughton, S. J., M. Oesterheld, D. A. Frank, and K. J. Williams. 1989. Ecosystem-level patterns of primary productivity and herbivory in terrestrial habitats. Nature 341:142– 144.

Mejia, F. H., A. K. Fremier, J. R. Benjamin, J. R. Bellmore, A. Z. Grimm, G. A. Watson, and M. Newsom. 2019. Stream metabolism increases with drainage area and peaks asynchronously across a stream network. Aquatic Sciences 81:9.

Moore, J. W., J. D. Yeakel, D. Peard, J. Lough, and M. Beere. 2014. Life-history diversity and its importance to population stability and persistence of a migratory fish: steelhead in two large North American watersheds. Journal of Animal Ecology 83:1035–1046.

Mueller, T., and W. F. Fagan. 2008. Search and navigation in dynamic environments - from individual behaviors to population distributions. Oikos 117:654–664.

Munson, B. H., J. H. McCormick, and H. L. Collins. 1980. Influence of Thermal Challenge on Conditioned Feeding Forays of Juvenile Rainbow Trout. Transactions of the American Fisheries Society 109:116–121.

Nakano, S., K. D. Fausch, and S. Kitano. 1999. Flexible niche partitioning via a foraging mode shift: a proposed mechanism for coexistence in stream-dwelling charrs. Journal of Animal Ecology 68:1079–1092.

Nakano, S., and M. Murakami. 2001. Reciprocal subsidies: Dynamic interdependence between terrestrial and aquatic food webs. Proceedings of the National Academy of Sciences 98:166–170.

Naman, S. M., J. S. Rosenfeld, P. M. Kiffney, and J. S. Richardson. 2018. The energetic consequences of habitat structure for forest stream salmonids. Journal of Animal Ecology 87:1383–1394.

Naman, S. M., S. M. White, J. R. Bellmore, P. A. McHugh, M. J. Kaylor, C. V. Baxter, R. J. Danehy, R. J. Naiman, and A. L. Puls. 2022. Food web perspectives and methods for riverine fish conservation. WIREs Water 9.

Nielsen, J. L. 1992. Microhabitat-Specific Foraging Behavior, Diet, and Growth of Juvenile Coho Salmon. Transactions of the American Fisheries Society 121:617–634.

Oldham, T., B. Nowak, M. Hvas, and F. Oppedal. 2019. Metabolic and functional impacts of hypoxia vary with size in Atlantic salmon. Comparative Biochemistry and Physiology Part A: Molecular & Integrative Physiology 231:30–38.

Patten, M. A., and J. F. Kelly. 2010. Habitat selection and the perceptual trap. Ecological Applications 20:2148–2156.

Pépino, M., K. Goyer, and P. Magnan. 2015. Heat transfer in fish: are short excursions between habitats a thermoregulatory behaviour to exploit resources in an unfavourable thermal environment? Journal of Experimental Biology:jeb.126466.

Piccolo, J. J., B. M. Frank, and J. W. Hayes. 2014. Food and space revisited: The role of drift-feeding theory in predicting the distribution, growth, and abundance of stream salmonids. Environmental Biology of Fishes 97:475–488.

Poff, N. L., and A. D. Huryn. 1998. Multi-scale determinants of secondary production in Atlantic salmon (*Salmo salar*) streams. Canadian Journal of Fisheries and Aquatic Sciences 55:201–217.

Power, M. E. 1984. Habitat Quality and the Distribution of Algae-Grazing Catfish in a Panamanian Stream. The Journal of Animal Ecology 53:357.

Power, M.E. and W.E. Rainey. 2000. Food webs and resource sheds: Towards spatially delimiting trophic interactions. In Ecological Consequences of Habitat Heterogeneity, ed. M.J. Hutchings, E.A. John and A.J.A. Stewart. Oxford, UK: Blackwell Scientific, pp. 291–314.

Power, M.E., Dietrich W.E., O’Sullivan K. 1998. Experimentation, Observation, and Inference in River and Watershed Investigations. Experimental Ecology 6:113–132

Price, N., L. Lopez, A. E. Platts, and J. R. Lasky. 2020. In the presence of population structure: From genomics to candidate genes underlying local adaptation. Ecology and Evolution 10:1889–1904.

Quaempts, E. J., K. L. Jones, S. J. O’Daniel, T. J. Beechie, and G. C. Poole. 2018. Aligning environmental management with ecosystem resilience: A First Foods example from the Confederated Tribes of the Umatilla Indian Reservation, Oregon, USA. Ecology and Society 23:29.

Quinn, T. P. 2018. The behavior and ecology of Pacific salmon and trout. Second edition. University of Washington PressJ; In association with American Fisheries Society, SeattleJ: Bethesda, Maryland.

Rahel, F. J., and J. W. Nutzman. 1994. Foraging in a Lethal Environment: Fish Predation in Hypoxic Waters of a Stratified Lake. Ecology 75:1246–1253.

Railsback SF, Harvey BC, Hayse JW, LaGory KE. 2005. Tests of theory for diel variation in salmonid feeding activity and habitat use. Ecology. 86:947–959.

Rossi, G. J., M. E. Power, S. M. Carlson, and T. E. Grantham. 2022. Seasonal growth potential of *Oncorhynchus mykiss* in streams with contrasting prey phenology and streamflow. Ecosphere 13 (9) e4211.

Rossi, G. J., M. E. Power, S. Pneh, J. R. Neuswanger, and T. J. Caldwell. 2021. Foraging modes and movements of *Oncorhynchus mykiss* as flow and invertebrate drift recede in a California stream. Canadian Journal of Fisheries and Aquatic Sciences 78:1045–1056.

Rossi, G.. J. 2020. Food, Phenology, and Flow—How Prey Phenology and Streamflow Dynamics Affect the Behavior, Ecology, and Recovery of Pacific Salmon. UC Berkeley. ProQuest ID: 19607. Retrieved from https://www.etdadmin.com/etdadmin/files/135/745674_pdf_876894_64127B12-9F6B-11EA-A155-7B0A2F6F8D95.pdf

Rüegg, J., C. C. Conn, E. P. Anderson, T. J. Battin, E. S. Bernhardt, M. Boix Canadell, S. M. Bonjour, J. D. Hosen, N. S. Marzolf, and C. B. Yackulic. 2021. Thinking like a consumer: Linking aquatic basal metabolism and consumer dynamics. Limnology and Oceanography Letters 6:1–17.

Ruff, C. P., D. E. Schindler, J. B. Armstrong, K. T. Bentley, G. T. Brooks, G. W. Holtgrieve, M. T. McGlauflin, C. E. Torgersen, and J. E. Seeb. 2011. Temperature-associated population diversity in salmon confers benefits to mobile consumers. Ecology 92:2073–2084.

Ryser, R., M. R. Hirt, J. Häussler, D. Gravel, and U. Brose. 2021. Landscape heterogeneity buffers biodiversity of simulated meta-food-webs under global change through rescue and drainage effects. Nature Communications 12:4716.

Schindler, D. E., R. Hilborn, B. Chasco, C. P. Boatright, T. P. Quinn, L. A. Rogers, and M. S. Webster. 2010. Population diversity and the portfolio effect in an exploited species. Nature 465:609–612.

Schlosser, I. J. 1991. Stream Fish Ecology: A Landscape Perspective. BioScience 41:704–712.

Sims, D. W., M. J. Witt, A. J. Richardson, E. J. Southall, and J. D. Metcalfe. 2006. Encounter success of free-ranging marine predator movements across a dynamic prey landscape. Proceedings of the Royal Society B: Biological Sciences 273:1195–1201.

Sinclair, A. R. E., and R. Beyers. 2021. A place like no other: discovering the secrets of Serengeti. Princeton University Press, Princeton.

Sommer, T. R., M. L. Nobriga, W. C. Harrell, W. Batham, and W. J. Kimmerer. 2001. Floodplain rearing of juvenile chinook salmon: evidence of enhanced growth and survival. Canadian Journal of Fisheries and Aquatic Sciences 58:325–333.

Stanford, J. A., M. S. Lorang, and F. R. Hauer. 2005. The shifting habitat mosaic of river ecosystems. SIL Proceedings, 1922-2010 29:123–136.

Sturrock, A. M., M. Ogaz, K. Neal, N. J. Corline, R. Peek, D. Myers, S. Schluep, M. Levinson, R. C. Johnson, and C. A. Jeffres. 2022. Floodplain trophic subsidies in a modified river network: managed foodscapes of the future? Landscape Ecology 37:2991–3009.

Twining, C. W., T. P. Parmar, M. Mathieu-Resuge, M. J. Kainz, J. R. Shipley, and D. Martin-Creuzburg. 2021. Use of Fatty Acids From Aquatic Prey Varies With Foraging Strategy. Frontiers in Ecology and Evolution 9:735350.

van der Graaf, A. J., Stahl, J., Klimkowska, A., Bakker, J. P., Drent, R. H. 2006. Surfing on a green wave: How plant growth drives spring migration in the Barnacle Goose Branta leucopsis. Ardea, 94(3), 567 577.

Warren, C. E., and G. E. Davis. 1967. Laboratory studies on the feeding, bioenergetics and growth of fish, p. 175-214. In S. D. Gerking [ed.] The biological basis of freshwater fish production. Blackwell Scientific Publications, Oxford. 510 p.

White, S.M., G.R. Giannico, and H.W. Li. 2014. A ‘Behaviorscape’ Perspective on Stream Fish Ecology and Conservation: Linking Fish Behavior to Riverscapes. Wiley Interdisciplinary Reviews: Water 1, no. 4: 385–400. 10.1002/wat2.1033.

Whitney, E. J., J. R. Bellmore, J. R. Benjamin, C. E. Jordan, J. B. Dunham, M. Newsom, and M. Nahorniak. 2020. Beyond sticks and stones: Integrating physical and ecological conditions into watershed restoration assessments using a food web modeling approach. Food Webs 25:e00160.

Wiens, John A. 2002. Riverine Landscapes: Taking Landscape Ecology into the Water. Freshwater Biology 47:501–15.

Wipfli, M. S., and C. V. Baxter. 2010. Linking Ecosystems, Food Webs, and Fish Production: Subsidies in Salmonid Watersheds. Fisheries 35:373–387.

Woodson, L., B. Wells, P. Weber, R. MacFarlane, G. Whitman, and R. Johnson. 2013. Size, growth, and origin-dependent mortality of juvenile Chinook salmon Oncorhynchus tshawytscha during early ocean residence. Marine Ecology Progress Series 487:163–175.

Wootton, J. T., M. S. Parker, and M. E. Power. 1996. Effects of Disturbance on River Food Webs. Science 273:1558–1561.

Závorka, L., A. Crespel, N. J. Dawson, M. Papatheodoulou, S. S. Killen, and M. J. Kainz. 2021. Climate changeJinduced deprivation of dietary essential fatty acids can reduce growth and mitochondrial efficiency of wild juvenile salmon. Functional Ecology 35:1960–1971.

